# Effect of G_4_C_2_ Repeat Expansions on the Motion of Lysosomes Inside Neurites

**DOI:** 10.1101/2021.10.29.466472

**Authors:** Maria Mytiliniou, Joeri A. J. Wondergem, Marleen Feliksik, Thomas Schmidt, Doris Heinrich

**Affiliations:** Leiden Institute of Physics, Huygens-Kamerlingh Onnes Laboratory, Leiden University, Leiden, the Netherlands; Institute for Bioprocessing and Analytical Measurement Techniques, Rosenhof, 37308 Heilbad Heiligenstadt, Germany; Faculty for Mathematics and Natural Sciences, Technische Universität Ilmenau, Ilmenau, Germany; Fraunhofer Institute for Silicate Research ISC, Würzburg, Germany

## Abstract

The G_4_C_2_ hexanucleotide repeat expansion in the c9orf72 locus is one among a plethora of mutations associated with amyotrophic lateral sclerosis. It accounts for the majority of disease cases. The exact processes underlying the pathology of this mutation remain elusive, yet recent evidence suggests a mechanism that disrupts axonal trafficking. Here, we used a neuronal cell line with and without the G_4_C_2_ repeats, and implemented time-resolved local mean squared displacement analysis to characterize the motion of lysosomes inside neurites. Neurites were either aligned along chemically patterned lines, or oriented randomly on the substrate. We confirmed that in the presence of the G_4_C_2_ repeats, lysosome motion was affected. Lysosomes had a smaller reach exhibited lower velocity, especially inside aligned neurites. At the same time they became more active with increasing length of the G_4_C_2_ repeats when the neurites were randomly oriented. The duration of diffusive and super-diffusive lysosome transport remained unaffected for both neurite geometries and for all lengths of the repeats, but the displacement and velocity was decreased on varying the repeat number and neurite geometry. Lastly, the ratio of anterograde/retrograde/neutral trajectories was affected disparately for the two neurite geometries. Our observations support the hypothesis that impaired axonal trafficking emerges in the presence of the G_4_C_2_ hexanucleotide repeat expansion.

## Introduction

Cellular metabolic activities are intricately regulated and any imbalances can contribute to the pathology of various human diseases. The autophagy pathway is one such metabolic process, responsible for the degradation of intracellular components, for instance aggregated proteins and foreign bodies, and entails the fine tuning of a plethora of processes, among which, lysosome transport [1]. Amyotrophic lateral sclerosis (ALS) is a devastating, terminal neurodegenerative disease, with diverse genetic causes, a multitude of which, share one common characteristic: a connection with the autophagy pathway [2].

A mutation in chromosome 9, open reading frame 72 (c9orf72) associated with ALS was discovered and reported by two independent research groups in 2011 [3, 4], and has been drawing significant attention since, as it accounts for more ALS cases than any other known ALS-associated mutation [5]. The malfunction is caused by a G_4_C_2_ repeat expansion in a non-coding region of the c9orf72 locus. Non-affected, healthy individuals have n=2-19 (G_4_C_2_)_*n*_ units, whereas ALS patients have hundreds, or even thousands of the repeat units [6]. Higher numbers of the hexanucleotide repeat expansion (HRE) have been associated with increased cellular toxicity [7], earlier disease onset [8–10], or more acute pathological phenotype [10–13] however, patient cases, carrying as little as n=24-38 (G_4_C_2_)_*n*_ repeat units have been reported [10, 14–18].

The (G_4_C_2_)_*n*_ expansion is transcribed into RNA, in all six reading frames of the sense and anti-sense direction, forming stable G-quadruplexes or hairpins, which aggregate to RNA foci [6, 19–21]. In addition, all RNA products are translated, via non-ATG initiated (RAN) translation, into non-functional dipeptide-repeat (DPR) proteins: glycine-alanine (GA), glycine-arginine (GR), proline-alanine (PA), proline-arginine (PR) and glycine-proline (GP, generated from both the sense and anti-sense reading frames) [22–29].

Numerous studies have focused on understanding the underlying pathogenic mechanisms, their potential association with the RNA foci, the accumulated DPR proteins, or the loss-of-function of the normal gene product, and the development of possible therapeutics (reviewed in [10]). To comprehend the underlying pathogenicity, it is essential to be aware of the normal function of the gene product. Thus, various studies which focused on this question reveal that the c9orf72 protein is implicated, among others, with the autophagy-lysosome pathway and with lysosome homeostasis (reviewed in [30, 31]). Hence, it is not surprising that research on the c9orf72 repeat expansion mutation and its associated RNA and DPR protein products demonstrate evidence of disrupted vesicle trafficking [32] and autophagy [33–41]. In fact, two recent studies have reported disrupted axonal trafficking of lysosomes [42] and mitochondria [43] in the presence of the c9orf72 mutation.

Along the same lines, and aiming to shed more light on the association between the HRE products and defective axonal transport, we used our previously established experimental setup of neuron-like, differentiated PC12 cells, on a chemically-patterned substrate [44, 45]. In that study, we have shown that we can identify and quantitatively characterize the effect of a sucrose-induced perturbation on lysosomal motion, which differs with the neurite geometry.

Here, we confirmed an effect on lysosomal motion, inside cells carrying the G_4_C_2_ repeat expansion products. Interestingly, we found that the diffusive and super-diffusive transport modes of lysosomes inside aligned neurites, and the sub-diffusive transport parts inside randomly oriented neurites, were influenced the most. Moreover, in differentiated PC12 cells producing the (G_4_C_2_)_20_ or (G_4_C_2_)_39_ RNA and DPR proteins, lysosomes traversed smaller distances and exhibited lower velocity, more notably inside aligned neurites. The average *α* exponent value and the percentage of super-diffusive motion manifested an increase with increasing repeats number, when the neurites were randomly oriented, suggesting more enhanced transport. In addition, our results showed that the duration of diffusive and super-diffusive motion remained unaffected for both neurite geometries, and numbers of repeats. On the contrary, the displacement and velocity of all transport modes exhibited a decrease, varying with the repeat number and neurite geometry. Lastly, we demonstrated an effect on the ratio of anterograde/retrograde/neutral trajectories caused by the (G_4_C_2_)_20_ and (G_4_C_2_)_39_ products, which varied for the two neurite geometries. Interestingly, the instantaneous velocity and the repeats-associated decrease were the same for both directions in each neurite geometry. In conclusion, our findings underpin the hypothesis that the HRE gives rise to ALS pathology and associates with defects in axonal transport of lysosomes.

## Results

### Dynamic analysis of lysosomal trajectories inside neurites of varying geometry

Cell shape can influence intracellular processes, such as the transport of organelles, vesicles and molecules. Here, we analyzed the motion of lysosomes inside neurites of two different geometries. In the first configuration, the neurites were guided to grow along chemically-patterned, with attractant protein, lines, whereas in the second configuration, they were allowed to adopt any orientation and geometry on the two-dimensional surface. The generation of both neurites geometries has been described in detail previously [44]. Representative volume-view images of two differentiated PC12 cells and their neurites adopting the respective geometries are shown in Fig. 1.A and B. The pattern used to promote neurite guidance is demonstrated in the fluorescent image of inset Fig. 1.a, and indicated by the dashed white lines in Fig. 1.A. The thick side-stripe of the ladder-shape pattern allowed attachment of the cell body, and the dissecting 2*μ*m-wide lines guided the neurite outgrowths.

**Figure 1:**
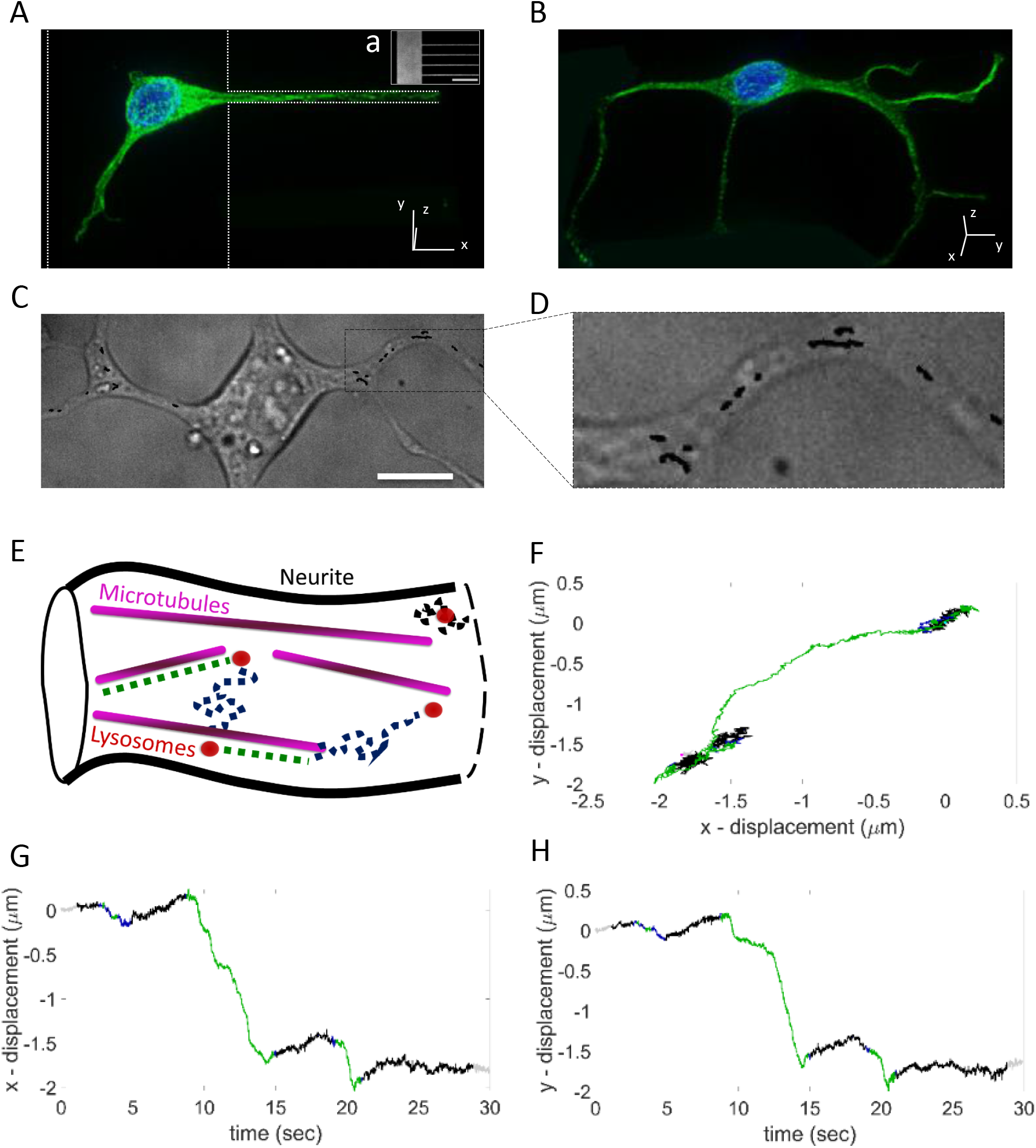
Neurites configurations and transport modes of lysosomes. (A) Volume-view image of a differentiated PC12 cell with its neurite aligning along the 2*μ*m-wide patterned line. The white dashed line indicates the underlying protein pattern. Green color corresponds to alpha-tubulin and blue to the cell nucleus. Inset (a) shows the fluorescently-labeled patterned substrate, used to promote one-dimensional neurite guidance. Scale bar 50*μ*m. (B) Volume-view image of a differentiated PC12 cell with the neurites adopting random orientation on the flat substrate. Green color corresponds to alpha-tubulin and blue to the cell nucleus. (C) Bright-field image of a differentiated PC12 cell, overlayed with lysosomes trajectories (black lines). Scale bar 10*μ*m. (D) Close-up of the area indicated by the black square in (C). (E) Schematic of the three lysosome transport modes, which were distinguished by local MSD analysis: sub-diffusion (black dashed line), diffusion (blue dashed line) and super-diffusion, including active transport (green dashed line). (F) Displacement along the x- and y- axis of a lysosome trajectory, color-coded for the three transport modes: sub-diffusion (black), diffusion (blue) and super-diffusion (green). The motion mode of each trajectory data point was determined based on the *α* exponent value, after fitting the local MSD. *α* < 0.9 corresponds to sub-diffusive motion, 0.9 ≤ *α* ≤ 1.1 corresponds to diffusive motion and *α* > 1.1 to super-diffusive motion. (G) and (H) Displacement along the x- and y- axis respectively, as a function of time, of the trajectory shown in (F).

Trajectories of lysosomes inside neurites were monitored and their coordinates as a function of time were recorded. Fig. 1.C and D shows an example bright-field image of a differentiated cell, overlayed with recorded trajectories of lysosomes. To quantify the lysosomal dynamics, local MSD (lMSD) analysis was used [46], resulting in time-resolved characterization of the trajectory into sub-diffusive (*α* < 0.9), diffusive (0.9 ≤ *α* ≤ 1.1) or super-diffusive (*α* > 1.1) motion. We considered a window of ~ 1*sec* before and after each data point to calculate the lMSD, which we subsequently fit for lag times 0-555 msec, to determine *α*. Using this analysis we have previously identified and quantitatively characterized a sucrose-induced perturbation in the cellular environment that affects lysosomal motion, and we have verified the fact that neurite geometry has a distinct impact on organelle transport [44].

The sketch in Fig. 1.E illustrates the three motion modes, and a lysosomal trajectory exhibiting all three modes is plotted in Fig. 1.F. Usually, diffusive or sub-diffusive states are intersected by rapid, super-diffusive translocation, part of which entails motor-mediated transport. It is very common also that no active transport is observed within a trajectory, and the lysosome is undergoing only passive (sub-) diffusion. In addition, part of the super-diffusive motion might not be motor-protein-mediated (for which *α* ≈ 2), but instead, a result of thrust from the fibrilar network or from cytoplasmic flow.

The three motion modes alternate stochastically with each other. Super-diffusive motion corresponds to large displacements in short time, with high directionality, as can be seen in Fig. 1.G and H. Diffusive and sub-diffusive motion result in smaller net displacements and a high degree of randomness.

### Neuronal cell model carrying (G_4_C_2_)_20_ or (G_4_C_2_)_39_

The (G_4_C_2_)_*n*_ hexanucleotide repeat expansion at the c9orf72 locus has been studied extensively during the last decade, as it accounts for more ALS cases than any other known ALS-associated mutation. The normal function of the protein is now known to be associated with the autophagy-lysosome pathway and there have been two recent reports indicating the c9orf72 mutation as a cause of disrupted axonal mitochondria and lysosome transport [42, 43].

To characterize the motion of lysosomes in the presence of the repeat expansion, we transfected PC12 cells with plasmids carrying (G_4_C_2_)_20_ or (G_4_C_2_)_39_, preceded by a start codon aligned such as to result in the production of (GR)_20_ or (GA)_39_, respectively. Multiple studies have shown that poly-GA is the most abundant of the five DPR protein species in patients [47, 48], and cells [28, 49, 50]. In addition, this DPR protein is very aggregation-prone and triggers pathological phenotypes in a zebrafish model [51]. Likewise, arginine-rich DPR proteins, such as the poly-GR, have shown higher expression than other DPR protein species [28] and correlation with pathological phenotypes in patient cells [52], fly model [53], chick embryo [54], *C.elegans* [55], and zebrafish [56]

We confirmed the presence of repeats-carrying transcribed RNA in the transfected cells, by quantifying the smFISH signal (Fig. S1). A five-fold increase was observed in the signal of cells transfected with (G_4_C_2_)_20_, as compared with the control case (the auto-fluorescence of the cells), and an almost two-fold increase in comparison with the non-specific binding signal of the FISH probes. The increase of the FISH fluorescence intensity in cells transfected with (G_4_C_2_)_39_ was less than with (G_4_C_2_)_20_, as compared to both respective controls, but that could be attributed to the RNA being more aggregation-prone, thus hindering binding of the FISH probes.

### In the presence of G_4_C_2_ repeats, the mobility and velocity of lysosomes decreases, most notably inside aligned neurites

To gain a quantitative insight on the effect of the presence of the G_4_C_2_ products on lysosomal motion, we first looked at the maximum Euclidean displacement observed in a trajectory and the average instantaneous velocity, for each experimental condition. As displayed in Fig. 2 (black bars), the maximum displacement per trajectory was, on average, the same for lysosomes inside aligned and randomly oriented neurites, extending to slightly less than 0.5*μm*. The instantaneous velocity was slightly increased for lysosomes inside aligned neurites, ≊ 0.24*μm/sec*, versus ≊ 0.2*μm/sec* inside randomly oriented neurites.

**Figure 2:**
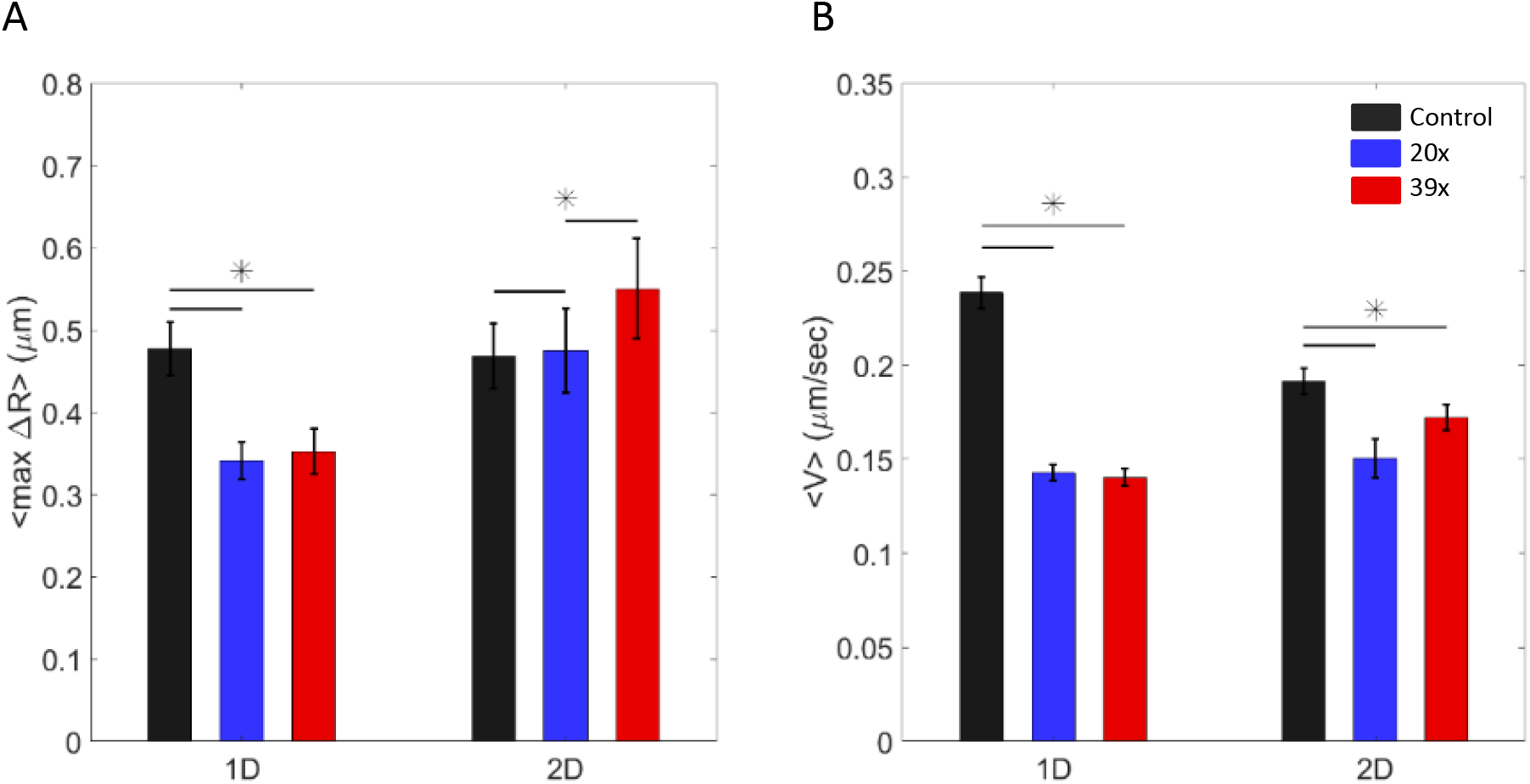
Alterations in the maximum displacement and instantaneous velocity of lysosomes in the presence of the G_4_**C**_2_ repeats. (A) Mean value of the maximum Euclidean displacement (B) Average instantaneous velocity. Values were calculated per trajectory and averaged over all trajectories of lysosomes per experimental condition (inside aligned (1D) or randomly oriented (2D) neurites, of differentiated PC12 cells without (control, black) or with the G_4_C_2_ repeat expansion ((G_4_C_2_)_20_, blue and (G_4_C_2_)_39_, red, respectively)). * indicates significant difference (*p* 0.05), as determined using the Wilcoxon ranksum test. Error bars indicate the standard error of the mean.

In the presence of (G_4_C_2_)_20_ and (G_4_C_2_)_39_ (blue and red bars, respectively, in Fig. 2), lysosomes moving inside aligned neurites exhibited the same significant decrease in both their maximum displacement (≊ 0.15*μm*), and the average instantaneous velocity (≊ 0.1*μm/sec*), as compared to the control. When the neurites were randomly oriented, the instantaneous velocity exhibited a significant, although smaller (< 0.05*μm/sec*) decrease than the respective drop observed in the case of the aligned neurites, when the cells carried the G_4_C_2_ repeat expansion. However, the maximum displacement was not significantly affected in the case of lysosomal motion inside randomly oriented neurites, in the presence of (G_4_C_2_)_20_ and (G_4_C_2_)_39_. Instead, it appeared to be marginally increased.

Thus, so far, it appears that 1D neurite alignment gave rise to slightly faster lysosomal motion, though not larger displacements. Moreover, this neurite geometry associates with greater decrease in both the instantaneous velocity and average maximum displacement in the presence of the (G_4_C_2_)_20_ or (G_4_C_2_)_39_ RNA and DPR proteins products, as compared to neurites that are randomly oriented.

### The proportion among the lysosomal motion modes is unchanged by the presence or length of the G_4_C_2_ repeat expansion

Next, we wondered whether the presence and size of the G_4_C_2_ repeat expansion has a distinct effect on the type of motion and the proportion among the transport modes, as a function of the neurite geometry. Thus, we looked into the average value of the *α* exponent per trajectory, and what percentage of the motion per trajectory is super-diffusive.

As can be seen in Fig. 3.A, the overall mean *α* value per trajectory is notably less than one (black bars), indicating that lysosomes spent a high fraction of their motion in the sub-diffusive mode. When the neurites were aligned, the observed variance among the mean *α* values of lysosomal trajectories inside cells carrying the repeat expansion and the control case, is not significant in the case of the (G_4_C_2_)_20_ (blue bars), whereas it is significant but still smaller than 0.1 for the (G_4_C_2_)_39_ (red bars). In the case where the neurites adopted a random geometry, the mean *α* value of the respective lysosomal trajectories increased significantly only in the presence of the (G_4_C_2_)_39_ repeat expansion, resulting in an *α* value approximately double the corresponding control value.

**Figure 3:**
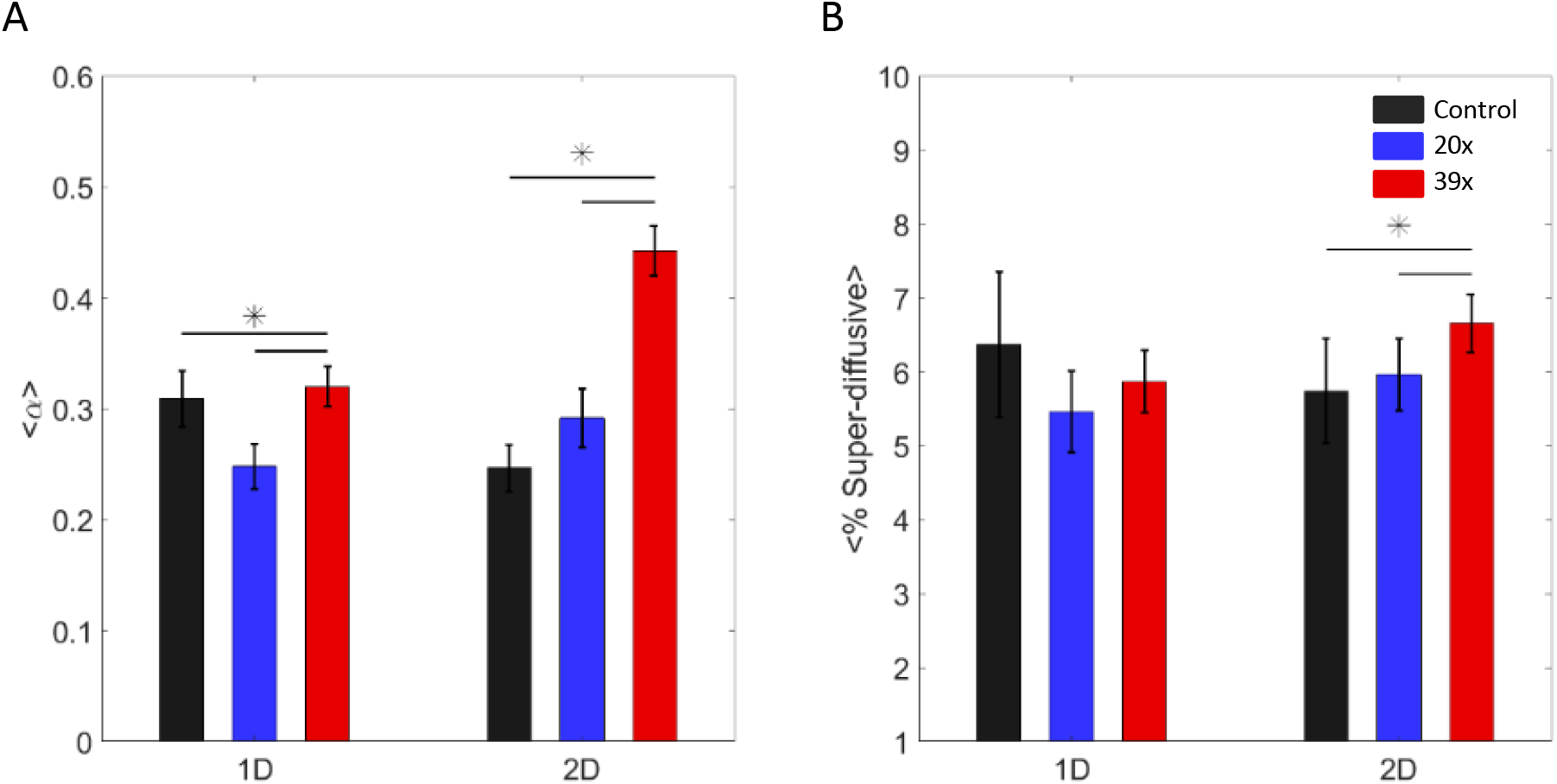
Average *α* exponent value and percentage of super-diffusive motion of lysosomal trajectories inside neurites of differentiated PC12 cells without or with the G_4_C_2_ repeats. (A) Mean value of the *α* exponent. (B) Average proportion of super-diffusive data points. Values were calculated per trajectory and averaged over all trajectories of lysosomes per experimental condition (inside aligned (1D) or randomly oriented (2D) neurites, of differentiated PC12 cells without (control, black) or with the G_4_C_2_ repeat expansion ((G_4_C_2_)_20_, blue and (G_4_C_2_)_39_, red, respectively)). * indicates significant difference (*p* ≤ 0.05), as determined using the Wilcoxon ranksum test. Error bars indicate the standard error of the mean.

The percentage of super-diffusive motion, displayed in Fig. 3.B, exhibited no significant difference for trajectories inside aligned neurites and significant, but in the order of only 1% increase as compared to the control, for trajectories inside randomly oriented neurites, only in the presence of the (G_4_C_2_)_39_ repeat expansion.

All in all, these results imply a general effect on the cellular environment, and specifically on lysosomal motion, caused by the presence of the (G_4_C_2_)_39_ and none for the (G_4_C_2_)_20_ RNA and DPR proteins products, when the neurites are allowed to adopt a random geometry.

### The G_4_C_2_ repeat expansion associates with decreased lysosomal displacement and instantaneous velocity for all transport modes

Thereafter, we sought for a deeper insight into the effect of the presence of the (G_4_C_2_)_20_ and (G_4_C_2_)_39_ products inside the cell, in combination with the two distinct neurite geometries, on each of the three lysosome transport modes. To this end, we broke down every single trajectory into parts of each transport mode, as determined by the *α* exponent value resulting from the lMSD analysis. Then, we analyzed collectively all the trajectory parts for each experimental condition. The corresponding distributions of the duration, maximum displacement, and instantaneous velocity of the three motion modes for each experimental condition are displayed in Fig. 4, 5 and 6, respectively. The corresponding mean values of the distributions are summarized in Tables S1, S2 and S3.

**Figure 4:**
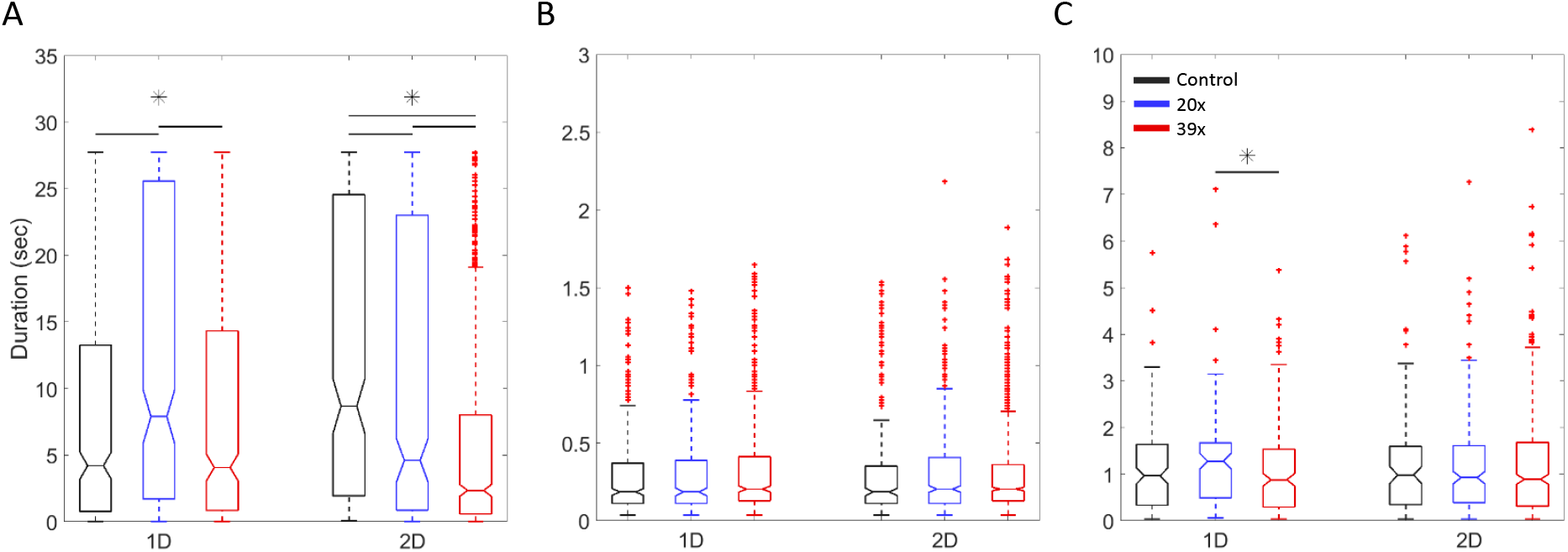
Duration of trajectory modes. Mean duration of (A) sub-diffusive, (B) diffusive and (C) super-diffusive trajectory modes, averaged over all the respective trajectory parts of lysosomes for each condition (aligned (1D) and randomly oriented (2D) neurites, of differentiated PC12 cells without (control, black) and with the G_4_C_2_ repeat expansion ((G_4_C_2_)_20_, blue and (G_4_C_2_)_39_, red)). * indicates significant difference (*p* ≤ 0.05), as determined using the Wilcoxon ranksum test.

**Figure 5:**
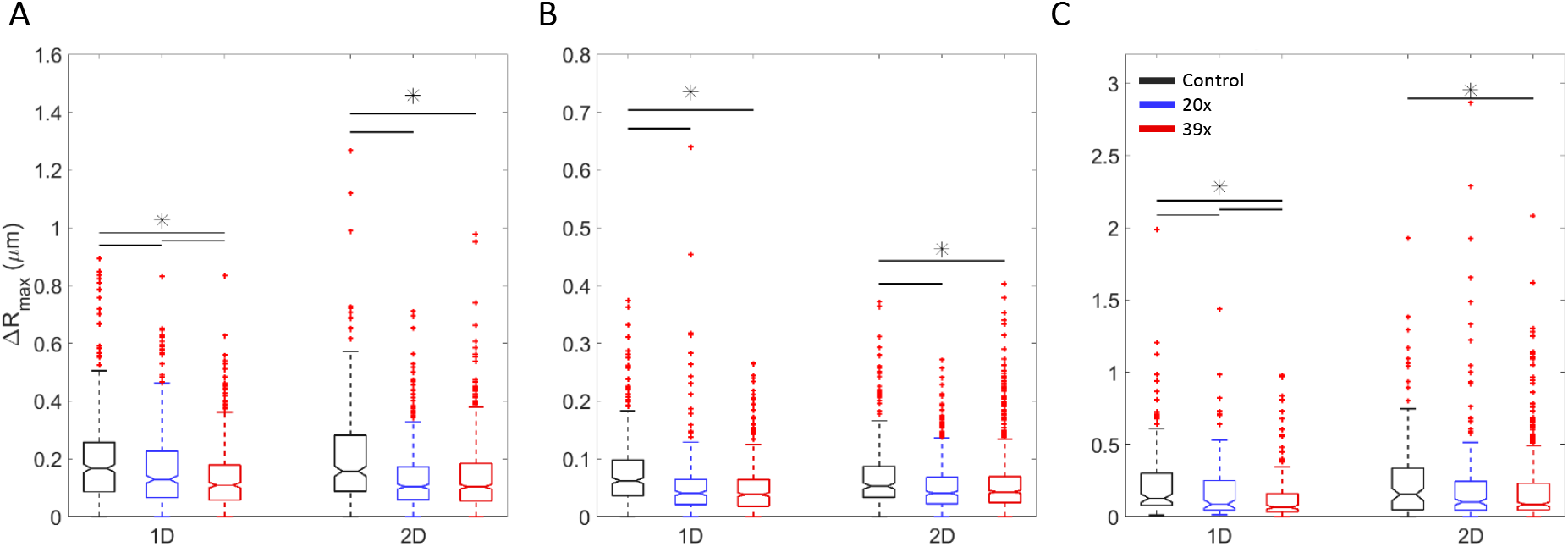
Maximum displacement of trajectory modes. Maximum Euclidean displacement observed during (A) sub-diffusive, (B) diffusive and (C) super-diffusive trajectory modes, averaged over all the respective trajectory parts of lysosomes for each condition (aligned (1D) and randomly oriented (2D) neurites, of differentiated PC12 cells without (control, black) and with the G_4_C_2_ repeat expansion ((G_4_C_2_)_20_, blue and (G_4_C_2_)_39_, red)). * indicates significant difference (*p* ≤ 0.05), as determined using the Wilcoxon ranksum test.

**Figure 6:**
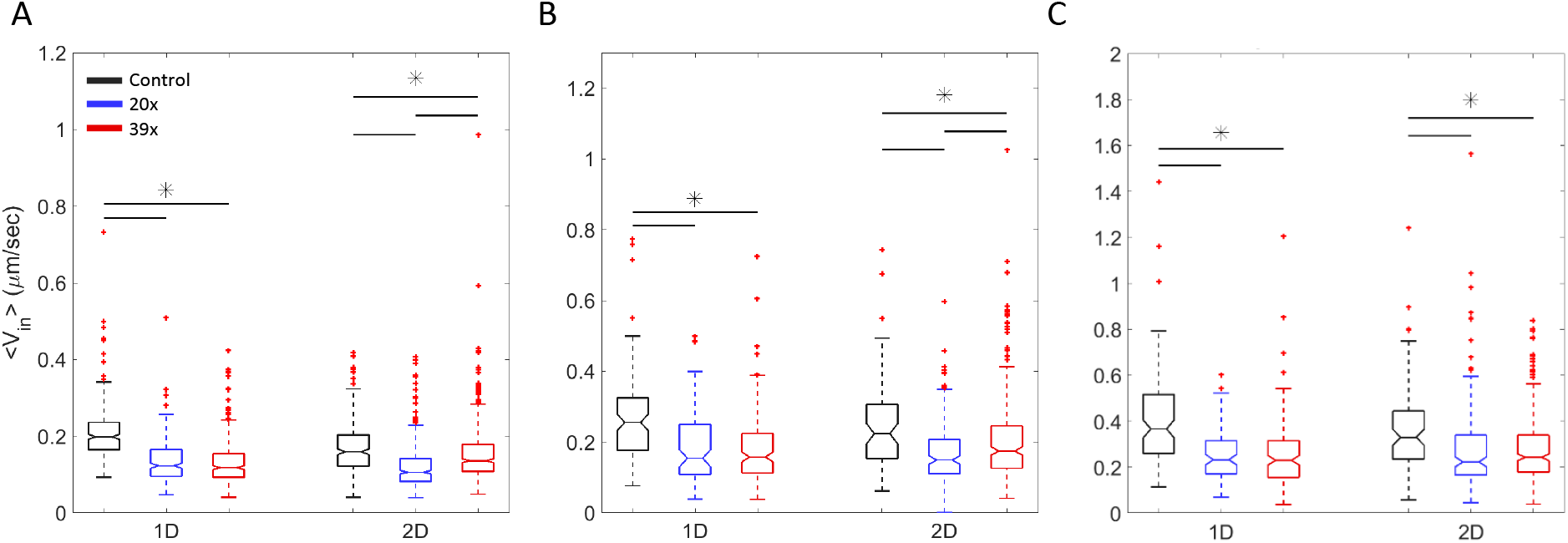
Average instantaneous velocity of trajectory modes. Average instantaneous velocity during (A) sub-diffusive, (B) diffusive and (C) super-diffusive trajectory modes, averaged over all the respective trajectory parts of lysosomes for each condition (aligned (1D) and randomly oriented (2D) neurites, of differentiated PC12 cells without (control, black) and with the G_4_C_2_ repeat expansion ((G_4_C_2_)_20_, blue and (G_4_C_2_)_39_, red)). * indicates significant difference (*p* ≤ 0.05), as determined using the Wilcoxon ranksum test.

The duration of diffusive and super-diffusive transport modes appears unaffected by the presence of the repeat expansion products or the neurite geometry. On average, diffusive transport states lasted for ≈ 300*msec*, and super-diffusive states for ≈ 1.2*sec*. Sub-diffusive modes duration exhibited a statistically significant decrease of approximately one second in the presence of (G_4_C_2_)_20_, with regards to the value in the cells without the repeat expansion, inside randomly oriented neurites. The respective decrease for cells transfected with (G_4_C_2_)_39_, was approximately 6 seconds, resulting in a value that is almost half the state duration of the control condition. The drop attenuated when the neurites were aligned; the sub-diffusive states duration remained the same in the presence of (G_4_C_2_)_39_, and increased by almost 4 seconds in the presence of (G_4_C_2_)_20_. Overall, sub-diffusive transport modes showed the longest duration (between 6 and 12 seconds). Thus, the observed trajectories (with a maximum duration of 30 seconds) entailed a large proportion of sub-diffusive motion, intersected by brief (super-) diffusive phases.

Concerning the maximum Euclidean displacement observed during each motion state, our results show that it decreased inside cells carrying the G_4_C_2_ repeat expansion, for all transport modes. Specifically, inside aligned neurites of cells transfected with (G_4_C_2_)_20_, the maximum displacement was decreased by 15%, 29.5% and 27.5%, as compared to the cells without the repeats, in the case of the sub-diffusive, diffusive and super-diffusive transport modes, respectively. The corresponding reduction was 30% for sub-diffusive, 37% for diffusive and 44% for super-diffusive lysosomal transport modes inside aligned neurites of cells transfected with (G_4_C_2_)_39_. Inside neurites of a random geometry, the maximum displacement decrease was 38% for sub-diffusive, 27% for diffusive and 16% for super-diffusive modes of lysosomes inside cells transfected with (G_4_C_2_)_20_ and 35% for sub-diffusive, 24% for diffusive and 23% for super-diffusive lysosomal transport modes of cells transfected with (G_4_C_2_)_39_, as compared to the respective values of motion modes inside cells without the repeats. Lastly, in the control case, the mean value of the calculated maximum displacement per transport mode exhibited negligible difference between the two neurite geometries.

The average instantaneous velocity of lysosomal trajectory modes exhibited similar values for both neurite geometries (~ 0.20, ~ 0.26 and ~ 0.38*μm/sec* for the sub-diffusive, diffusive and super-diffusive modes, respectively). A significant decrease was observed for all transport modes, in both neurite geometries and in the presence of either of the repeats lengths, as compared to the respective values for cells without the G_4_C_2_ repeats. Interestingly, when neurites were aligned, the decrease per mode was similar for both geometries; 36-37% for sub- and super- diffusive modes and 31-34% for diffusive modes. On the other hand, when neurites were randomly oriented, the corresponding decrease observed was 28%, 32% and 22% in cells transfected with the 20x repeats and 10%, 18% and 24% in cells transfected with the 39x repeats, for sub-diffusive, diffusive and super-diffusive transport modes, respectively.

### Fitting parameters of transport modes MSD curves confirm the effect of the G_4_C_2_ repeat expansion

Subsequently, we calculated the MSDs of the sub-diffusive, diffusive and super-diffusive parts of lysosomal trajectories. We fitted the sub-diffusive MSD curves with a power law, commonly used to describe anomalous diffusion (eq. 6). Likewise, we used the Brownian motion model (eq. 7) to fit the diffusive MSDs. Lastly, to fit the super-diffusive MSD curves we implemented the model of Brownian motion with drift (eq. 7) [57]. The MSD curves with the corresponding fits are shown in Fig. 7 and the resulting fitting values are summarized in Table S4.

**Figure 7:**
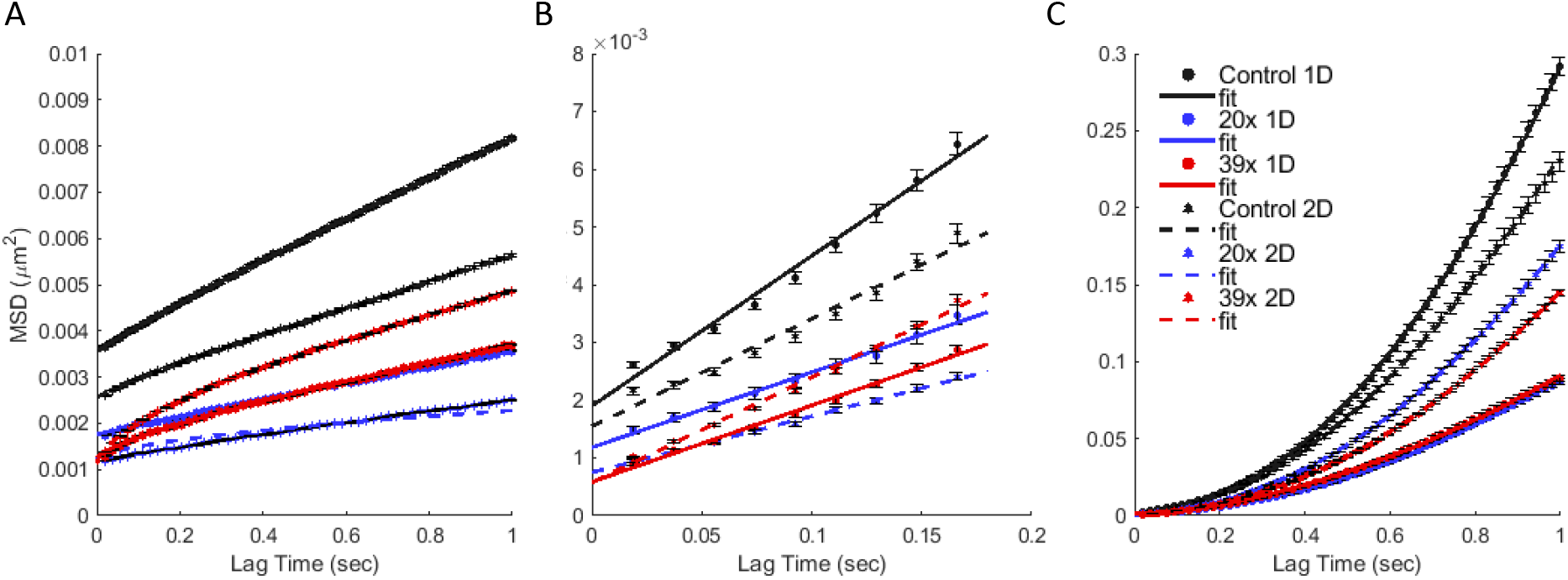
MSD plots of motion modes. MSD plots and respective fits of (A) sub-diffusive, (B) diffusive and (C) super-diffusive trajectory modes, averaged over all the respective trajectory parts of lysosomes for each condition: inside aligned (1D) and randomly oriented (2D) neurites, of differentiated PC12 cells without (control, black) and with the G_4_C_2_ repeat expansion ((G_4_C_2_)_20_, blue and (G_4_C_2_)_39_, red, respectively). The MSD curves were fitted using eq. 6-8 and the resulting values of the fitting parameters are summarized in Table S4.

A decrease was observed in the *α* exponent of the sub-diffusive motion for cells transfected with (G_4_C_2_)_20_ or (G_4_C_2_)_39_, as compared to the control, more prominent when the neurites adopt a random geometry as compared to when they are aligned. In the latter case, the decrease was only observed for the cells carrying the (G_4_C_2_)_39_.

Inside aligned neurites, the diffusion coefficient of the diffusive lysosomal transport modes was reduced to almost half the respective control value, in the presence of both (G_4_C_2_)_20_ or (G_4_C_2_)_39_ repeats products (~ 0.0033*μm*^2^/*sec* versus ~ 0.0065*μm*^2^/*sec* the control value). Similar decrease exhibited the drift velocity of the super-diffusive trajectory parts (~ 0.27 *μm/sec* versus ~ 0.54*μm/sec* the control value).

Contrary to that, and concerning lysosomes inside randomly oriented neurites, the diffusion coefficient of the diffusive transport modes was reduced to half, in the presence of (G_4_C_2_)_20_ (~ 0.0024*μm*^2^/*sec*), but remained the same in the presence of (G_4_C_2_)_39_ (~ 0.0046*μm*^2^/*sec*), as compared to the respective control value (~ 0.0047*μm*^2^/*sec*). The corresponding drift velocity of the super-diffusive trajectory parts was smaller in the presence of the repeats (~ 0.37 − 0.41*μm/sec*) as compared to the control case (~ 0.46*μm/sec*), but the drop was much less than the one observed for aligned neurites.

Thus, these results confirm an effect of the G_4_C_2_ repeat expansion products, more prominent for the diffusive and super-diffusive transport modes of lysosomes inside aligned neurites, and for the sub-diffusive transport parts inside randomly oriented neurites.

### Motion statistics do not differentiate for anterograde and retrograde motion inside aligned neurites

Lastly, we investigated whether the presence of the G_4_C_2_ repeat expansion products had a different effect on lysosomal motion towards (retrograde) or away from (anterograde) the cell body. To this end, we categorized accordingly all trajectories that exhibited super-diffusive motion, based on their net displacement (end point versus start point). In multiple occasions, the net displacement was zero or perpendicular to the axis pointing towards or away from the cell body, thus in this case the trajectory was labeled as neutral.

The proportion of each type of trajectory for every experimental condition is summarized in Fig. 8.A. It turns out that inside aligned neurites, less lysosomes moved away from the cell body and more trajectories exhibited a neutral net displacement, when the cells carried the repeat expansion, as compared to the control case. Surprisingly, the exact opposite was observed for lysosomes inside neurites of a random geometry.

**Figure 8:**
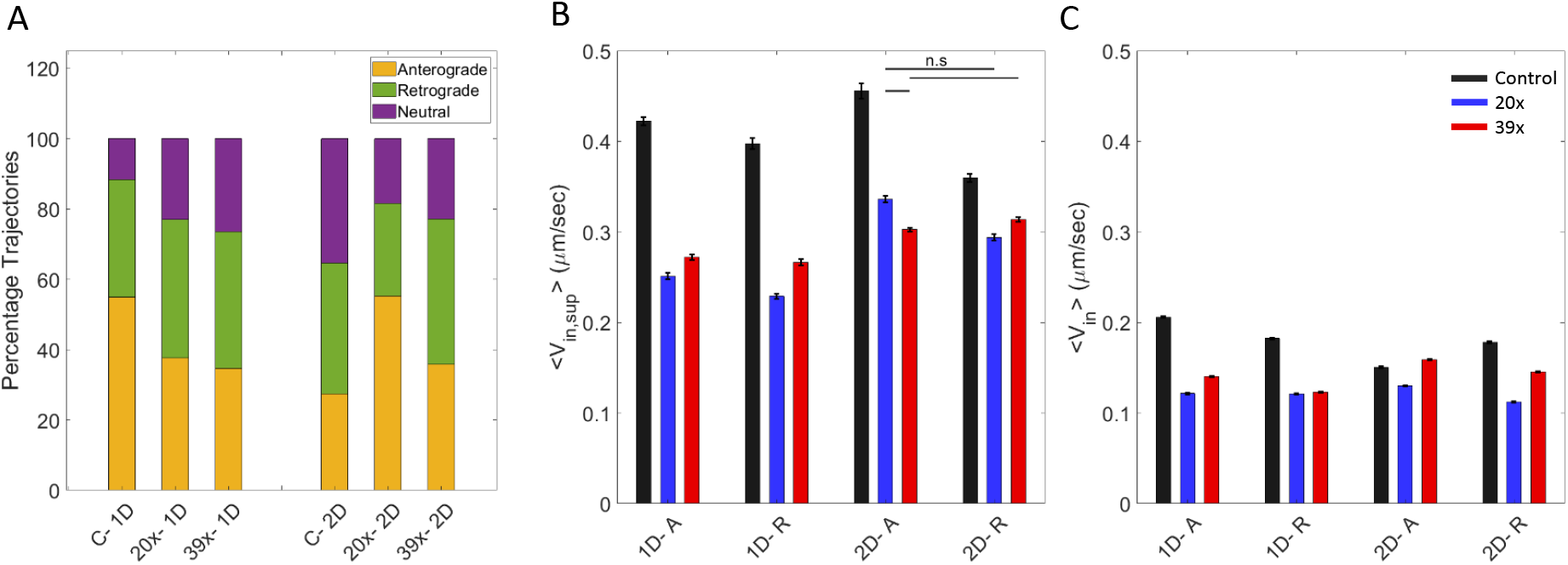
Statistics on anterograde and retrograde super-diffusive trajectories. (A) Percentage of lysosomal trajectories exhibiting super-diffusive motion, distinguished according to their net displacement either towards (retrograde) or away from (anterograde) the cell body for each condition: inside aligned (1D) and randomly oriented (2D) neurites, of differentiated PC12 cells without (control) and with the G_4_C_2_ repeat expansion ((G_4_C_2_)_20_ and (G_4_C_2_)_39_, respectively). In all cases, trajectories categorized as neutral didn’t exhibit a net displacement towards either direction. (B) Average instantaneous velocity of data points with *α* > 1.1 and (C) average instantaneous velocity of data points with *α* ≤ 1.1, from lysosomal trajectories with super-diffusive motion, exhibiting anterograde (A) or retrograde (R) net displacement inside aligned (1D) and randomly oriented (2D) neurites of differentiated PC12 cells. The mean values of the distributions were compared using the Wilcoxon ranksum test and they exhibited significant difference (*p* ≤ 0.05), unless indicated otherwise.

In addition, as shown in Fig. 8.B and C, the mean instantaneous velocities of both super-diffusive and diffusive data points of these trajectories exhibited the same decrease regardless the direction of motion (anterograde or retrograde) and the size of the repeats, for lysosomes inside aligned neurites. The same trend is maintained for lysosomes inside randomly oriented neurites, but with a smaller decrease.

To summarize, these results indicate that the ratio of anterograde/retrograde/neutral trajectories changes differently for the two neurite geometries in the presence of the (G_4_C_2_) repeats. On the contrary, the decrease observed in the instantaneous velocity when the cells carry the repeats, is the same for anterograde/retrograde motion in each respective neurite geometry.

## Discussion

In order to aid the progress of a therapeutic development for the devastating disease of ALS, a thorough understanding of the underlying pathogenic mechanisms is necessary. Here, we set our focus on quantifying the effect of the G_4_C_2_ repeats, the ALS-associated mutation on the c9orf72 locus, on lysosomal motion. The protein encoded by the c9orf72 is implicated with the autophagy-lysosome pathway [31, 36, 58–67], and lysosome homeostasis [58, 68], while recent reports indicate the c9orf72 mutation as a cause for deficient axonal transport of mitochondria [42] and lysosomes [43]. Using a model system of differentiated PC12 cells transfected with plasmids containing either (G_4_C_2_)_20_ or (G_4_C_2_)_39_, we strived to verify these recent findings, by monitoring and analyzing the motion of lysosomes inside the cells’ neurites.

For both repeat lengths investigated here ((G_4_C_2_)_20_ and (G_4_C_2_)_39_), patient cases have been identified and experimental models have been reported to recapitulate pathological hallmarks [10, 14–18]. Our results consolidate these findings, since we observed smaller displacements and velocities of lysosomes inside neurites of cells carrying either of the repeats lengths, as compared to the control case. Moreover, the decrease in the displacement was bigger for lysosomes inside cells carrying (G_4_C_2_)_39_ repeats as compared to (G_4_C_2_)_20_ repeats, supporting the claim of repeats length-dependent toxicity or disease onset [7–13]

Arginine-rich DPR proteins such as the poly-GR have been known for their high expression, as compared to other DPR proteins, and correlation with pathological phenotypes [28, 52, 53, 55, 56]. Fumagalli et al. recently contributed more knowledge on the connection between this DPR proteins species and axonal transport, and suggested a model where arginine-rich DPR proteins bind microtubules and concentrate in microtubule-associated foci, thereby disrupting the binding of motor proteins and subsequently efficient intracellular trafficking [43]. In agreement with this model, our results indicated decreased super-diffusive lysosomal displacement and velocity in the presence of (G_4_C_2_)_20_, translating into (GR)_20_. Notably, the velocity and diffusion coefficient values we obtained for lysosomes inside cells transfected with (G_4_C_2_)_20_ and (G_4_C_2_)_39_, are similar. This suggests that (G_4_C_2_)_20_ producing arginine-rich DPR proteins, affects lysosomal motion equally as the double repeats length (G_4_C_2_)_39_ producing alanine-rich DPR proteins. Thus, our finding further enhances the notion that arginine-rich products are highly correlated with pathological hallmarks, as compared to other DPR protein products.

Recently, Abo-Rady et al. reported decreased lysosomal displacement in axons of patient-derived motor neurons, but no significant alterations in the directionality of lysosomal trafficking [42]. In agreement with these results, we found decreased lysosomal displacement in all three motion modes, the same drop in the instantaneous velocity of anterograde and of retrograde trajectories, and only small variations in the ratio of anterograde/retrograde/neutral trajectories inside neurites of cells transfected with the (G_4_C_2_)_20_ and (G_4_C_2_)_39_.

Cell shape, greatly affected by the ECM architecture, can influence intracellular trafficking, among other processes [69–71]. In addition cell shape can associate with more acute pathological phenotype, as reported recently by Braun et al. who found increased severity of mechanically-induced tau pathology inside neurites which where aligned, as compared to neurites which were unaligned [72]. In the experiments presented here, except for investigating the effect of the ALS-associated HRE mutation on lysosomal trafficking inside neurites, we were in parallel looking into how the neurite geometry interferes with the motion. We have previously detected and quantitatively characterized a perturbation in the intracellular environment that affected lysosomal motion, while in parallel confirming a positive effect of neurite alignment on organelle motion, even in the presence of the perturbation [44]. Here, we verified a disturbance of lysosomal transport but we didn’t detect a consistently enhanced transport inside aligned neurites in the presence of the (G_4_C_2_)_20_ or (G_4_C_2_)_39_ RNA and DPR proteins. Instead, our results indicated larger decrease in the velocity and the displacement of lysosomes, particularly for their diffusive and super-diffusive transport modes, when those moved inside aligned neurites, suggesting an interesting interplay between the geometry of the cell and the effect of the repeat expansion on intracellular trafficking.

In summary, our results support the notion that defects in lysosome trafficking is a feature of ALS pathogenesis and, in the case of the c9orf72 mutation, a result of the accumulation of the hexanucleotide repeat expansion RNA and DPR proteins. It would be interesting, in a next step, to perform this quantitative characterization while discriminating between RNA foci or DPR protein accumulation, thus gaining more insight into the respective associated mechanisms. Of course, the possibility that other processes also contribute to the observed disrupted lysosomal motion is not excluded, and therefore the relative contribution of this deficit to the overall pathology remains elusive, however, our results support the notion that targeting the repair of axonal trafficking could constitute a potential therapeutic approach.

## Materials and Methods

### Laminin *μ*PIP

Plasma-initiated patterning of the substrate was performed as described previously [44]. Briefly, a mold was fabricated with a Nanoscribe Photonic professional GT 3D laser printer (Nanoscribe, Germany), with two-photon polymerization (2PP) of IP-S photoresist [73].

Poly dimethyl-siloxane (PDMS, Sylgard 184, Dow Corning, USA) was prepared by mixing the cross-linking agent with the elastomer base at a ratio of 1:10. The mixture was pipetted on the silicon mold and allowed to cross-link for 1 hour at 120°C. Subsequently, the hardened PDMS bearing the structure was peeled from the wafer.

The PDMS mask was placed on an ibidi dish (ibidi GMBH, *μ*-Dish, 35 mm high, polymer coverslip bottom), with the structure-bearing side adherent to the bottom of the dish and was exposed to air plasma for 6 min 20 sec at 100 Watts (Diener Electronic Femto Plasma system).

Subsequently, the PDMS mask was removed and the substrate was flooded with 0.1% Pluronic F127 (Sigma-Aldrich) diluted in Phosphate Buffer Saline (PBS) for 45 minutes at room temperature. The dish was then washed 3 times with PBS and once with RPMI (Gibco™) and subsequently incubated with 25*μ*g/ml Laminin (Sigma-Aldrich) diluted in RPMI, for 1 hour at 37°C. Prior to seeding cells, the substrate was washed 3 times with RPMI.

### Cell Culture

PC12 cells (CH3 BioSystems) were cultured in dishes coated with rat-tail Collagen (CH3 BioSystems), in RPMI-1640 with Glutamax (Gibco™), supplemented with 10% heat-inactivated Horse Serum (HS) (Sigma-Aldrich), 5% Fetal Calf Serum Heat Inactivated (FCS HI, Thermo Scientific) and 200*μ*g/mL pennicilin/streptomycin (PS). Media were refreshed three times per week, and the cells were split once per week at ratio 1:3-1:6. The cells were kept at 37°C and 5% CO_2_ in humidified atmosphere.

To induce differentiation, PC12 cells were seeded in ibidi uncoated dishes (ibidi GMBH, *μ*-Dish, 35 mm high) coated with Laminin (Sigma-Aldrich). Cells were seeded at a density of 30.000 cells/cm^2^in full media and after they had adhered, they were washed once with PBS and then the media were replaced with differentiation media consisting of Opti-MEMTM Reduced Serum Medium (Gibco™) supplemented with 0.5% Fetal Bovine Serum (Gibco™) and Nerve Growth Factor (NGF-2.5S Sigma-Aldrich) at final concentration of 100ng/ml. The differentiation media were refreshed three times per week.

Prior to imaging, the cells were incubated with 50-150nM Lysotracker (Invitrogen™) in RPMI for 30minutes.

Two plasmids, containing either 20 or 39 times the G_4_C_2_ sequence were created (GenScript) and amplified using NEB®Stable Competent *E.coli* bacteria (High Efficiency, New England BioLabs Inc.), according to the protocol for cloning DNA containing repeat elements (C3040, New England BioLabs Inc.). In both plasmids, the G_4_C_2_ repeat expansion was preceded by a start codon (ATG) and the reading frames were aligned such that the (G_4_C_2_)_20_ resulted in (GR)_20_ and the (G_4_C_2_)_39_ resulted in (GA)_39_ DPR proteins.

For transfection, ~ 5.7*x*10^5^ PC12 cells were seeded in each well of a Collagen-coated 6-well plate, and the next day they were transfected, using Lipofectamine LTX PLUS (Thermo-Fisher Scientific), with 2*μ*g of the (G_4_C_2_)_20_ or (G_4_C_2_)_39_ expression constructs per well. After a three-day incubation, Geneticin (G418, Thermo-Fisher Scientific) was added for selection of the transfected cells. After several days of selection, the cells were sparsely re-seeded on Collagen-coated 10-cm dishes to create single-cell clones, which were subsequently transferred to a Collagen-coated 96-well plate.

Successfully transfected clones were verified using single-molecule Fluorescent *in situ* Hybridization (smFISH). Briefly, the cells were fixed for 15 minutes with 4% PFA at room temperature and permeabilized overnight in 70% EtOH. Next, the custom-designed smFISH probes (LGC Biosearch Technologies, targeting 5’ GGCCCC GGCCCC GGCCCC GGCCCC 3’) were added at a concentration of 1:100 in hybridization buffer (100 mg/mL dextran sulfate, 25% formamide, 2X SSC, 1 mg/mL *E.coli* tRNA, 1 mM vanadyl ribonucleoside complex, 0.25 mg/mL BSA) and allowed to incubate for 16 hours at 30°C. Thereafter, the cells were washed twice with washing buffer (25% formamide, 2X SSC) containing DAPI (1 mg/mL, Sigma Aldrich), for 30 minutes at 30°C. All solutions were prepared using RNAse-free water. Lastly, the cells were mounted with Prolong Gold (Thermo-Fisher Scientific) and imaged two days later.

For immunofluorescence, differentiated PC12 cells were fixed with 4% PFA for 15 minutes at room temperature, permeabilized with 0.1% TritonX for 10 minutes and blocked with 2% BSA in PBS for 60 minutes. Subsequently the cells were incubated with the *α*-tubulin antibody (Alexa Fluor®488 anti-alpha Tubulin antibody, abcam) 1:150 in 0.1% BSA in PBS at 4°C for 18 hours. Lastly, the cells were incubated for 30 minutes at room temperature with Hoechst (Invitrogen) diluted to 1*μ*g/ml in 1% BSA in PBS and stored in PBS at 4°C until imaging.

### Optical Microscopy

Optical microscopy images were acquired with a Nikon Ti Eclipse inverted microscope (NIKON corporation, Japan) equipped with a Yokogawa CSU-X1 spinning disc unit (10,000rpm, Andor Technology Ltd., United Kingdom). The samples were imaged with a 100x objective (Nikon CFI Plan Apo Lamda, NA 1.45). Excitation at 405nm, 488nm and 647nm was achieved via an Agilent MLC400 monolithic laser combiner (Agilent Technologies, Netherlands). The excitation light was filtered by a custom-made Semrock quad-band dichroic mirror for excitation wavelengths 400-410, 486-491, 460-570, and 633-647nm. The emitted light was filtered using a Semrock quad-band fluorescence filter (TR-F440-521-607-700), which has specific transmission bands at 440±40nm, 521±21nm, 607±34nm and 700±45nm and by Semrock Brightline single band fluorescence filters at 447±60nm (TR-F447-060) and 525±60nm (TR-F525-030). Images were captured with an Andor iXon Ultra 897 High-speed EM-CCD camera. Image acquisition was automated using NisElements software (LIM, Czech Republic). Time-lapse images were acquired every 18 ms, for up to 30 seconds. During data acquisition, the cells were kept in a humidified atmosphere at 37°C and supplied with 5% CO_2_ via the use of a Tokai Hit stage incubator.

### Data Analysis

FISH signal quantification was performed with FIJI [74]. The integrated fluorescent intensity per cell nucleus was measured, after maximum intensity projection of the volume-view acquired images, and the values were averaged for a number of cells, N, per condition.

Lysosomes trajectories were tracked using the FIJI plugin, TrackMate [75], which returned the *x*– and *y*–coordinates of the center of the lysosomes, with a sub-pixel localization.

Further processing was performed using home-made Matlab algorithms. The *x*– and *y*–coordinates as a function of time for each trajectory were represented by a series of vectors at each time point *t*:

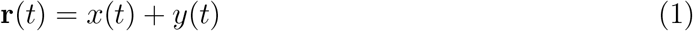

and the displacement Δ**r** at time *t* was calculated as follows:

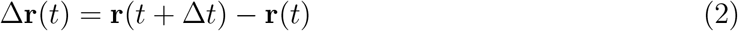

where Δ*t* is the inverse frame rate.

The instantaneous velocity *v*(*t*) was calculated using:

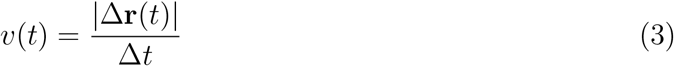

for Δ*t* = 314.5*msec* (17 frames).

The Mean Squared Displacement (MSD) for lag time *τ* = *k*Δ*t* was calculated according to:

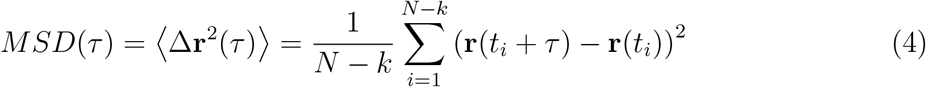

where *N* the number of data points in the trajectory and *k* = 1, 2, *…, N* − 1. The average MSD per condition is the average of the squared displacements of all lysosome trajectories for each lag time *τ*.

The local Mean Squared Displacement (lMSD) was calculated for each trajectory as described previously [44, 46]. Briefly, the MSD was calculated for each data point of the entire trajectory using a rolling window of 2.22 seconds (N=120, in eq. 4) and fitted for the interval 0-555ms (k=30 in eq. 4) with a power law:

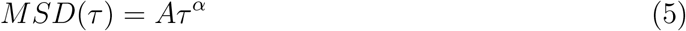

Tha alpha exponent as a function of time was subsequently used to partition the transport states as sub-diffusive for *α* < 0.9, diffusive for 0.9 ≤ *α* ≤ 1.1, or super-diffusive for *α* > 1.1.

In order to characterize more closely each type of motion, we analyzed collectively the respective trajectory parts, for each experimental condition (lysosomes inside neurites that were aligned or randomly oriented, of differentiated PC12 cells without the repeat expansion (control) or with 20x and 39x the G_4_C_2_ repeats).

The average MSD curve for each motion mode was calculated again according to eq. 4. The sub-diffusive trajectory modes MSD was fitted with the power law describing anomalous diffusion

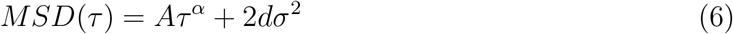

thereby obtaining the value of the anomalous *α* exponent (A is a constant). The MSD curve was fitted for all lag times (*τ* up to 30 sec).

The diffusive trajectory modes MSD was fitted using the equation describing Brownian motion

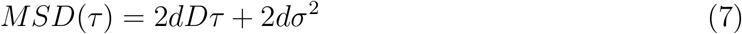

thus extracting the experimental value of the diffusion coefficient D. The MSD curve was fitted for lag times 1 to 10 (*τ* between 18.5 and 185 ms).

The super-diffusive trajectory modes MSD was fitted using the model of Brownian motion with drift [57].

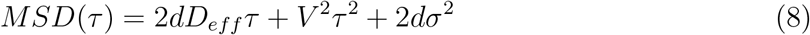

where *V_drift_*, the constant drift parameter models the velocity of the molecular motors. The MSD curve was fitted for lag times 1 to 55 (*τ* between 0.0185 and 1 sec).

In equations 6, 7 and 8 *σ* is the localization precision, *τ* the lag time and the parameter d refers to the dimensionality. d was set equal to 1 for the fit of the MSD along the x- or y-axis, and equal to 2 for the fit of the 2D- MSD curve.

The Jump Distance Distribution (JDD) was calculated according to the self-part of the van Hove correlation function [76]:

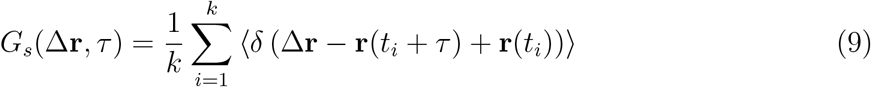

for displacements in both the *x*– and *y*– direction, and a specific lag time *τ*. *δ* here denotes the Dirac delta function in two dimensions and *k* = 1, 2, *…, N* − 1, with *N* the number of data points in the trajectory. The bin size was (arbitrarily) set to 1*μm* and the JDD was normalized into a probability density function.

The diffusive and super-diffusive trajectory parts for each experimental condition were used to calculate the respective JDD PDF, for lag times of 0.2405ms and 0.7585ms respectively. The PDFs were fitted to extract characteristic values of the motion, using the analytical expressions calculated in [77]. Particularly, the diffusive trajectory modes JDD PDF was fitted for the x- direction, using

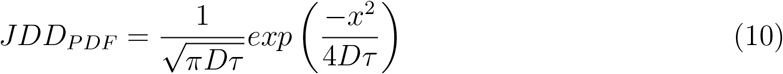

thereby estimating the experimental value of the diffusion coefficient D. Similarly, for the y-direction. The super-diffusive trajectory modes JDD PDF was fitted for the x- direction, using

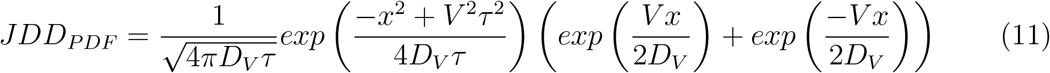

estimating the experimental value of the drift velocity *V_drift_* and effective diffusion coefficient *D_eff_*. Likewise for the y-direction.

## Acknowledgments

We thank Dr. Stefan Semrau (Leiden Institute of Physics, Leiden University, Leiden, the Netherlands) for the insightful discussions and assistance with smFISH. The authors would also like to acknowledge the Fraunhofer Gesellschaft for the Fraunhofer Attract “3DNanoCell”.

## Data Availability

Data is available on request from the corresponding author.

## Supporting Information

**Table S1:** Mean duration of sub-diffusive, diffusive and super-diffusive trajectory modes (in seconds). Values were averaged over all the respective trajectory parts of lysosomes for each condition (inside aligned (1D) and randomly oriented (2D) neurites, of differentiated PC12 cells without (control) and with the G_4_C_2_ repeat expansion ((G_4_C_2_)_20_ or (G_4_C_2_)_39_)). Errors were calculated using the standard error of the mean. The distributions of the values for each condition are displayed in Fig. 4.

|  | Sub-diffusive | Diffusive | Super-diffusive |
| --- | --- | --- | --- |
| 1D |  |  |  |
| Control | $8.13 \pm 0.50$ | $0.304 \pm 0.016$ | $1.13 \pm 0.084$ |
| 20x | $12.4 \pm 0.60$ | $0.313 \pm 0.018$ | $1.32 \pm 0.103$ |
| 39x | $8.59 \pm 0.46$ | $0.326 \pm 0.014$ | $1.07 \pm 0.067$ |
| 2D |  |  |  |
| Control | $11.8 \pm 0.60$ | $0.324 \pm 0.022$ | $1.27 \pm 0.116$ |
| 20x | $10.5 \pm 0.51$ | $0.324 \pm 0.014$ | $1.18 \pm 0.073$ |
| 39x | $6.28 \pm 0.31$ | $0.308 \pm 0.010$ | $1.21 \pm 0.057$ |

**Table S2:** Mean values of the maximum Euclidean displacement observed during sub-diffusive, diffusive and super-diffusive trajectory modes (in *μ*m). The values were averaged over all the respective trajectory parts of lysosomes for each condition (inside aligned (1D) and randomly oriented (2D) neurites, of differentiated PC12 cells without (control) and with the G_4_C_2_ repeat expansion ((G_4_C_2_)_20_ or (G_4_C_2_)_39_)). Errors were calculated using the standard error of the mean. The distributions of the values for each condition are displayed in Fig. 5.

|  | Sub-diffusive | Diffusive | Super-diffusive |
| --- | --- | --- | --- |
| 1D |  |  |  |
| Control | $0.200 \pm 0.008$ | $0.078 \pm 0.003$ | $0.251 \pm 0.025$ |
| 20x | $0.170 \pm 0.008$ | $0.055 \pm 0.004$ | $0.182 \pm 0.021$ |
| 39x | $0.139 \pm 0.005$ | $0.049 \pm 0.002$ | $0.140 \pm 0.013$ |
| 2D |  |  |  |
| Control | $0.210 \pm 0.010$ | $0.074 \pm 0.004$ | $0.265 \pm 0.030$ |
| 20x | $0.130 \pm 0.005$ | $0.054 \pm 0.002$ | $0.223 \pm 0.025$ |
| 39x | $0.136 \pm 0.005$ | $0.056 \pm 0.002$ | $0.203 \pm 0.018$ |

**Table S3:** Mean values of the instantaneous velocity during sub-diffusive, diffusive and super-diffusive trajectory modes (in*μ*m/sec). Values were averaged over all the respective trajectory parts of lysosomes for each condition (inside aligned (1D) and randomly oriented (2D) neurites, of differentiated PC12 cells without (control) and with the G_4_C_2_ repeat expansion ((G_4_C_2_)_20_ or (G_4_C_2_)_39_)). Errors were calculated using the standard error of the mean. The distributions of the values for each condition are displayed in Fig. 6.

|  | Sub-diffusive | Diffusive | Super-diffusive |
| --- | --- | --- | --- |
| 1D |  |  |  |
| Control | $0.208 \pm 0.004$ | $0.271 \pm 0.007$ | $0.395 \pm 0.019$ |
| 20x | $0.134 \pm 0.003$ | $0.186 \pm 0.006$ | $0.249 \pm 0.010$ |
| 39x | $0.132 \pm 0.003$ | $0.179 \pm 0.004$ | $0.250 \pm 0.010$ |
| 2D |  |  |  |
| Control | $0.170 \pm 0.004$ | $0.253 \pm 0.008$ | $0.362 \pm 0.018$ |
| 20x | $0.122 \pm 0.003$ | $0.173 \pm 0.004$ | $0.282 \pm 0.014$ |
| 39x | $0.153 \pm 0.003$ | $0.207 \pm 0.004$ | $0.276 \pm 0.007$ |

**Table S4:** Motion parameters resulting from fitting the sub-diffusive, diffusive and super-diffusive trajectory parts MSD curves (demonstrated in Fig. 7), using eq. 6, 7 and 8, respectively.

|  | Sub-diffusive | Diffusive | Super-diffusive |  |
| --- | --- | --- | --- | --- |
| | α- exponent | D ($\mu\text{m}^2/\text{sec}$) | $V_{drift}$ ($\mu\text{m}/\text{sec}$) | $D_{eff}$ ($\mu\text{m}^2/\text{sec}$) |
| 1D |  |  |  |  |
| Control | $0.92 \pm 0.01$ | $0.0065 \pm 0.0013$ | $0.538 \pm 0.008$ | $6E - 7 \pm 0.002$ |
| 20x | $0.97 \pm 0.04$ | $0.0033 \pm 0.0006$ | $0.278 \pm 0.007$ | $0.003 \pm 0.001$ |
| 39x | $0.74 \pm 0.05$ | $0.0033 \pm 0.0005$ | $0.258 \pm 0.009$ | $0.006 \pm 0.001$ |
| 2D |  |  |  |  |
| Control | $0.90 \pm 0.03$ | $0.0047 \pm 0.0015$ | $0.460 \pm 0.010$ | $0.005 \pm 0.002$ |
| 20x | $0.57 \pm 0.64$ | $0.0024 \pm 0.0003$ | $0.413 \pm 0.004$ | $0.001 \pm 0.001$ |
| 39x | $0.62 \pm 0.02$ | $0.0046 \pm 0.0007$ | $0.373 \pm 0.003$ | $0.001 \pm 0.001$ |
| | $\sigma$ ($\mu\text{m}$) | $\sigma$ ($\mu\text{m}$) | $\sigma$ ($\mu\text{m}$) | |
| 1D |  |  |  |  |
| Control | $0.030 \pm 0.003$ | $0.022 \pm 0.012$ | $0.022 \pm 0.022$ | |
| 20x | $0.021 \pm 0.003$ | $0.017 \pm 0.008$ | $0.008 \pm 0.015$ | |
| 39x | $0.018 \pm 0.004$ | $0.012 \pm 0.007$ | $0.000 \pm 0.014$ | |
| 2D |  |  |  |  |
| Control | $0.025 \pm 0.003$ | $0.020 \pm 0.012$ | $0.015 \pm 0.023$ | |
| 20x | $0.017 \pm 0.012$ | $0.014 \pm 0.006$ | $0.019 \pm 0.014$ | |
| 39x | $0.017 \pm 0.004$ | $0.012 \pm 0.009$ | $0.018 \pm 0.011$ | |

**Table S5:** Average instantaneous velocity during anterograde or retrograde motion of lysosomal trajectories. Values were calculated separately for data points with *α* > 1.1 (indicated as “sup”) or *α* ≤ 1.1, from trajectories with super-diffusive motion, exhibiting anterograde or retrograde net displacement inside aligned (1D) and randomly oriented (2D) neurites of differentiated PC12 cells without (control) and with the G_4_C_2_ repeat expansion ((G_4_C_2_)_20_ and (G_4_C_2_)_39_, respectively). Errors were represent the standard error of the mean. The respective distributions are shown in Fig. 8.B and C.

| | $V_{in,sup}$ ($\mu\text{m}/\text{sec}$) | | $V_{in}$ ($\mu\text{m}/\text{sec}$) | |
| --- | --- | --- | --- | --- |
|  | Anterograde | Retrograde | Anterograde | Retrograde |
| 1D |  |  |  |  |
| Control | $0.422 \pm 0.004$ | $0.398 \pm 0.006$ | $0.206 \pm 0.001$ | $0.182 \pm 0.001$ |
| 20x | $0.251 \pm 0.003$ | $0.229 \pm 0.003$ | $0.122 \pm 0.001$ | $0.121 \pm 0.001$ |
| 39x | $0.272 \pm 0.003$ | $0.267 \pm 0.003$ | $0.140 \pm 0.001$ | $0.123 \pm 0.001$ |
| 2D |  |  |  |  |
| Control | $0.456 \pm 0.008$ | $0.360 \pm 0.004$ | $0.150 \pm 0.001$ | $0.178 \pm 0.001$ |
| 20x | $0.336 \pm 0.003$ | $0.294 \pm 0.004$ | $0.130 \pm 0.001$ | $0.112 \pm 0.001$ |
| 39x | $0.303 \pm 0.002$ | $0.314 \pm 0.002$ | $0.159 \pm 0.001$ | $0.146 \pm 0.001$ |

**Table S6:**
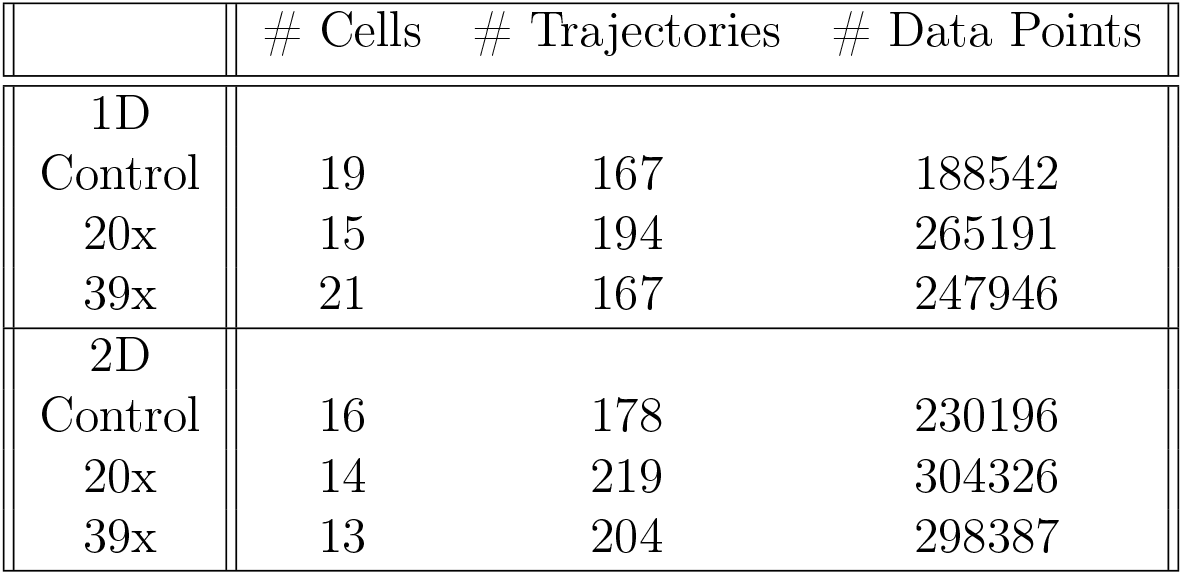
Number of cells imaged, number of total lysosomal trajectories tracked and number of total data points per experimental condition (differentiated PC12 cells without (control) or with the G_4_C_2_ repeats (20x and 39x), with their neurites aligned or randomly oriented, denoted by 1D and 2D, respectively).

|  | # Cells | # Trajectories | # Data Points |
| --- | --- | --- | --- |
| 1D |  |  |  |
| Control | 19 | 167 | 188542 |
| 20x | 15 | 194 | 265191 |
| 39x | 21 | 167 | 247946 |
| 2D |  |  |  |
| Control | 16 | 178 | 230196 |
| 20x | 14 | 219 | 304326 |
| 39x | 13 | 204 | 298387 |

**Table S7:**
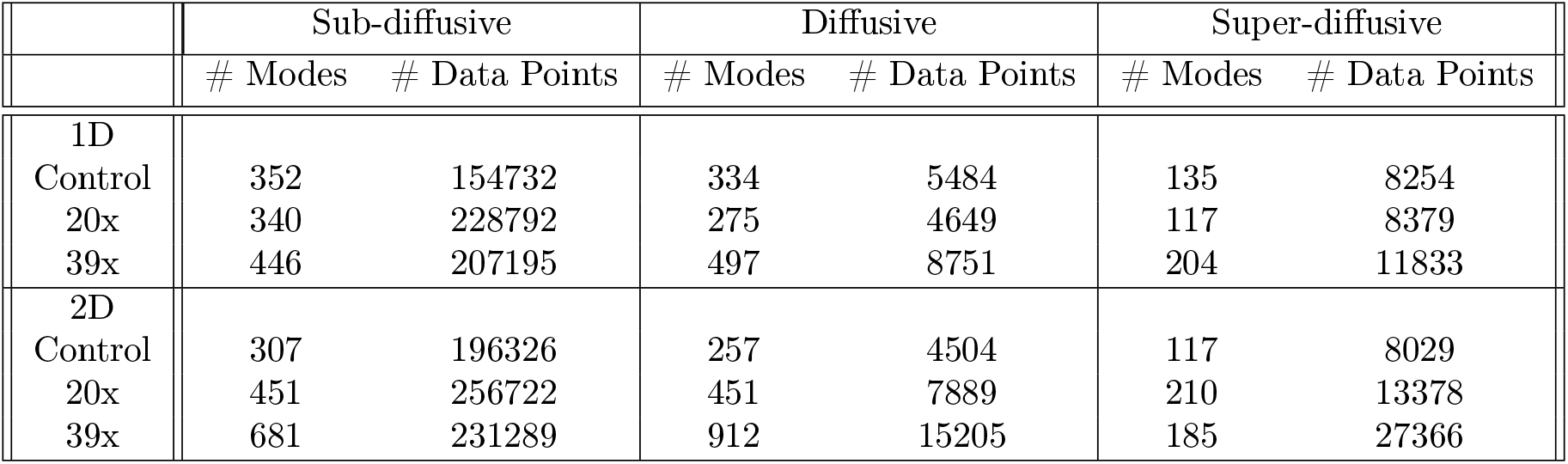
Number of lysosomal trajectories parts (modes) and number of total data points per mode, for each experimental condition (differentiated PC12 cells without (control) or with the G_4_C_2_ repeats (20x and 39x), with their neurites aligned or randomly oriented, denoted by 1D and 2D, respectively).

|  | Sub-diffusive |  | Diffusive |  | Super-diffusive |  |
| --- | --- | --- | --- | --- | --- | --- |
|  | # Modes | # Data Points | # Modes | # Data Points | # Modes | # Data Points |
| 1D |  |  |  |  |  |  |
| Control | 352 | 154732 | 334 | 5484 | 135 | 8254 |
| 20x | 340 | 228792 | 275 | 4649 | 117 | 8379 |
| 39x | 446 | 207195 | 497 | 8751 | 204 | 11833 |
| 2D |  |  |  |  |  |  |
| Control | 307 | 196326 | 257 | 4504 | 117 | 8029 |
| 20x | 451 | 256722 | 451 | 7889 | 210 | 13378 |
| 39x | 681 | 231289 | 912 | 15205 | 185 | 27366 |

**Table S8:** Motion parameters resulting from fitting the diffusive and super-diffusive trajectory parts x- and y- MSD curves (demonstrated in Fig. S2), using eq. 7 and 8.

| Motion mode | Diffusive |  | Super-diffusive |  |  |
| --- | --- | --- | --- | --- | --- |
| X- MSD | D ($\mu m^2/sec$) | $\sigma$ ($\mu m$) | $V_{drift}$ ($\mu m/sec$) | $D_{eff}$ ($\mu m^2/sec$) | $\sigma$ ($\mu m$) |
| 1D |  |  |  |  |  |
| Control | $0.0112 \pm 0.0025$ | $0.016 \pm 0.011$ | $0.525 \pm 0.002$ | $4E - 14 \pm 0.000$ | $0.012 \pm 0.017$ |
| 20x | $0.0057 \pm 0.0013$ | $0.014 \pm 0.008$ | $0.270 \pm 0.006$ | $0.0045 \pm 0.0017$ | $0.004 \pm 0.014$ |
| 39x | $0.0056 \pm 0.0011$ | $0.008 \pm 0.007$ | $0.264 \pm 0.007$ | $0.0094 \pm 0.0014$ | $7E - 6 \pm 0.000$ |
| 2D |  |  |  |  |  |
| Control | $0.0051 \pm 0.0015$ | $0.013 \pm 0.009$ | $0.351 \pm 0.006$ | $0.0038 \pm 0.0020$ | $0.014 \pm 0.015$ |
| 20x | $0.0018 \pm 0.0002$ | $0.010 \pm 0.003$ | $0.221 \pm 0.006$ | $0.0030 \pm 0.0013$ | $0.008 \pm 0.012$ |
| 39x | $0.0036 \pm 0.0005$ | $0.009 \pm 0.005$ | $0.175 \pm 0.004$ | $0.0032 \pm 0.0007$ | $0.006 \pm 0.009$ |
| Y- MSD |  |  |  |  |  |
| 1D |  |  |  |  |  |
| Control | $0.0018 \pm 0.0004$ | $0.015 \pm 0.004$ | $0.106 \pm 0.003$ | $0.0014 \pm 0.0003$ | $0.015 \pm 0.006$ |
| 20x | $0.0009 \pm 0.0002$ | $0.010 \pm 0.003$ | $0.069 \pm 0.006$ | $0.0010 \pm 0.0004$ | $0.009 \pm 0.007$ |
| 39x | $0.0010 \pm 0.0002$ | $0.008 \pm 0.003$ | $0.039 \pm 0.006$ | $0.0007 \pm 0.0002$ | $0.010 \pm 0.005$ |
| 2D |  |  |  |  |  |
| Control | $0.0042 \pm 0.0014$ | $0.014 \pm 0.008$ | $0.299 \pm 0.009$ | $0.0068 \pm 0.0029$ | $0.008 \pm 0.0018$ |
| 20x | $0.0031 \pm 0.0004$ | $0.009 \pm 0.005$ | $0.346 \pm 0.003$ | $0.00004 \pm 0.0012$ | $0.014 \pm 0.011$ |
| 39x | $0.0055 \pm 0.0009$ | $0.008 \pm 0.007$ | $0.328 \pm 0.003$ | $0.0001 \pm 0.0011$ | $0.014 \pm 0.011$ |

**Table S9:** Motion parameters resulting from fitting the diffusive and super-diffusive trajectory parts x- and y- JDD PDFs (demonstrated in Fig. S3), using eq. 10 and 11.

| Motion mode | Diffusive | Super-diffusive |  |
| --- | --- | --- | --- |
| X- JDD | D ($\mu m^2/sec$) | $V_{drift}$ ($\mu m/sec$) | $D_{eff}$ ($\mu m^2/sec$) |
| 1D |  |  |  |
| Control | $0.0112 \pm 0.0016$ | $0.388 \pm 0.060$ | $0.0186 \pm 0.0089$ |
| 20x | $0.0057 \pm 0.0007$ | $0.214 \pm 0.026$ | $0.0049 \pm 0.0019$ |
| 39x | $0.0050 \pm 0.0004$ | $0.225 \pm 0.033$ | $0.0065 \pm 0.0029$ |
| 2D |  |  |  |
| Control | $0.0054 \pm 0.0006$ | $0.205 \pm 0.024$ | $0.0050 \pm 0.0018$ |
| 20x | $0.0020 \pm 0.0002$ | $0.005 \pm 3147$ | $0.0180 \pm 11.8173$ |
| 39x | $0.0030 \pm 0.0003$ | $0.003 \pm 2498$ | $0.0113 \pm 5.7418$ |
| Y- JDD |  |  |  |
| 1D |  |  |  |
| Control | $0.0026 \pm 0.0003$ | $0.001 \pm 17931$ | $0.0043 \pm 12.7846$ |
| 20x | $0.0013 \pm 0.0001$ | $0.001 \pm 4958$ | $0.0019 \pm 3.7932$ |
| 39x | $0.0012 \pm 0.0001$ | $0.001 \pm 1155$ | $0.0013 \pm 1.0145$ |
| 2D |  |  |  |
| Control | $0.0042 \pm 0.0009$ | $0.007 \pm 5425$ | $0.0317 \pm 30.5804$ |
| 20x | $0.0024 \pm 0.0004$ | $0.174 \pm 0.076$ | $0.0150 \pm 0.0164$ |
| 39x | $0.0037 \pm 0.0005$ | $0.205 \pm 0.039$ | $0.0122 \pm 0.0075$ |

**Figure S1:**
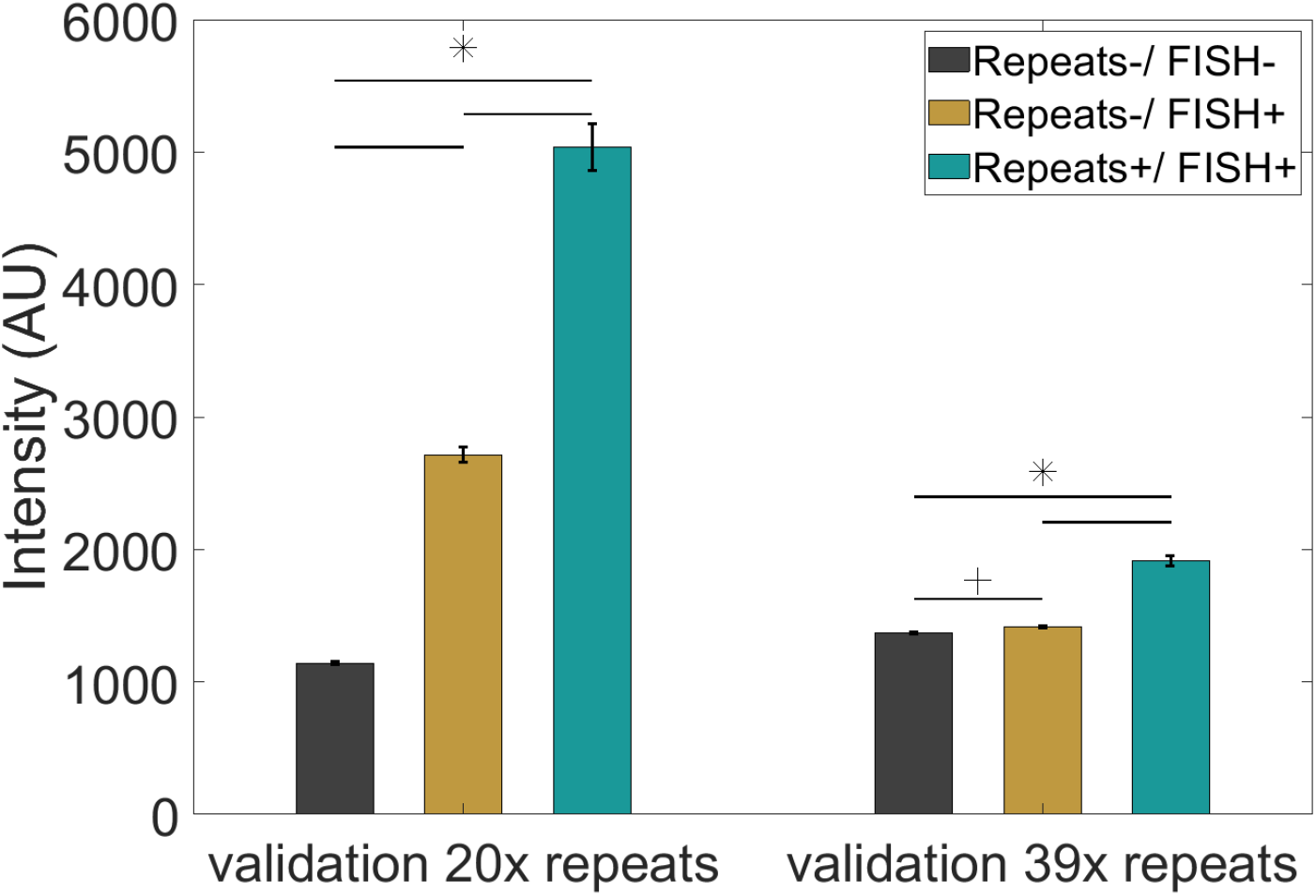
Verification of the presence of the (G_4_C_2_)_20_ or (G_4_C_2_)_39_ repeat expansion inside PC12 cells after transfection, and antibiotic selection. Validation was performed with FISH labeling against (G_4_C_2_)_4_ and subsequent quantification of the average nuclear fluorescent intensity per cell. Bars show the mean value ± SEM per condition: non-transfected cells, without FISH labeling (black- representing the cells’ intrinsic auto-fluorescence; N =55 for (G_4_C_2_)_20_ and N=64 cells for (G_4_C_2_)_39_ repeats validation), non-transfected cells, with FISH labeling (yellow- quantifying the non-specific binding of the FISH probes; N=88 for (G_4_C_2_)_20_ and N=145 cells for (G_4_C_2_)_39_ repeats validation), and cells transfected with either (G_4_C_2_)_20_ or (G_4_C_2_)_39_, with FISH labeling (green; N=100 for (G_4_C_2_)_20_ and N=166 cells for the (G_4_C_2_)_39_ validation). The mean values of the distributions were compared using the Wilcoxon ranksum test. + stands for *p* ≤ 0.05 and * for *p* < 0.001.

**Figure S2:**
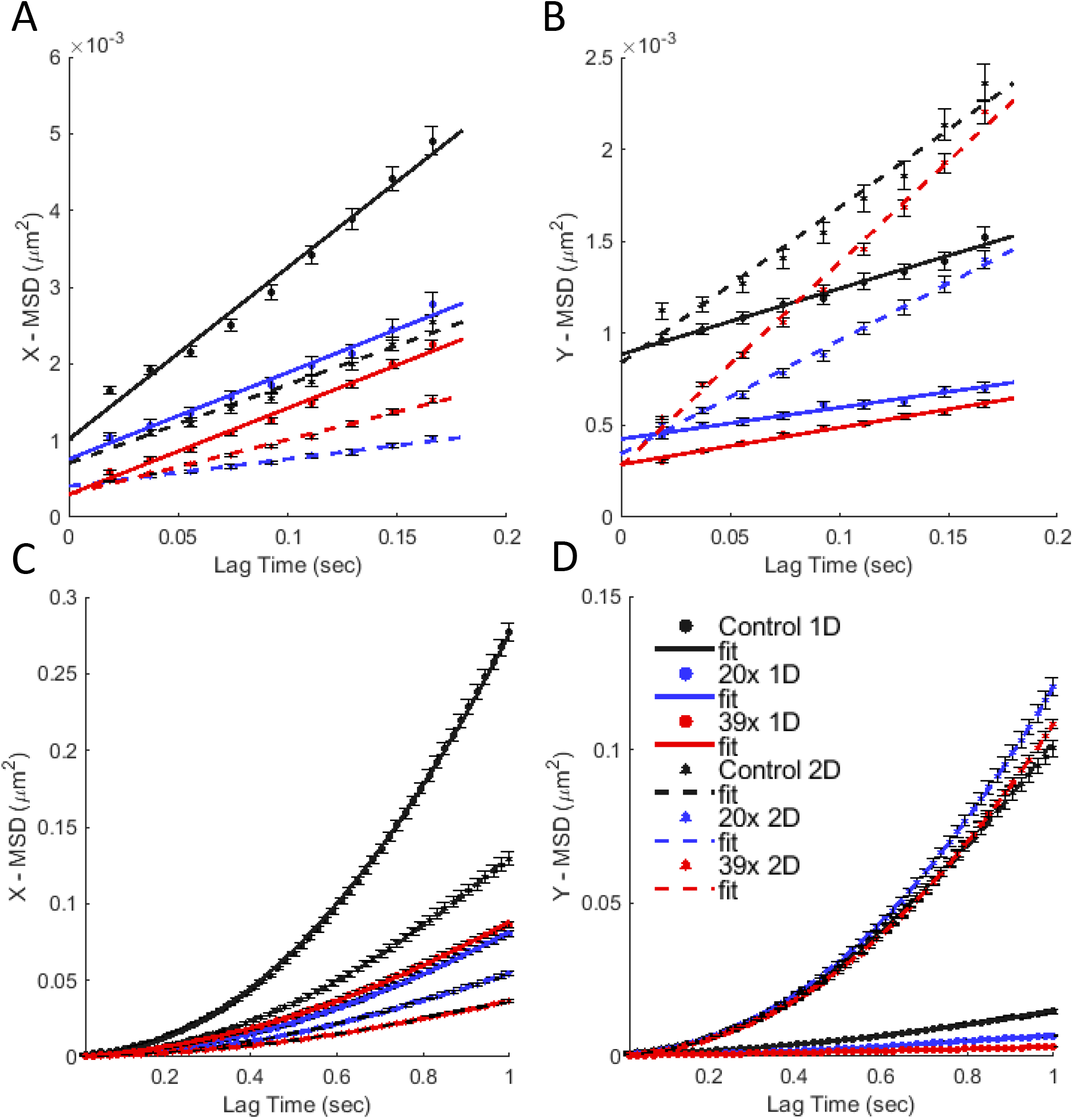
X- and Y- MSDs of diffusive and super-diffusive trajectory parts. MSDs calculated collectively for all diffusive trajectory parts along (A) the X- and (B) Y- axis. MSDs calculated collectively for all super-diffusive trajectory parts along (C) the X- and (D) Y-axis. Color coding indicates data of lysosomes inside aligned (1D) and randomly oriented (2D) neurites of differentiated PC12 cells without (control) and with the G_4_C_2_ repeat expansion (20x and 39x repeats, respectively). Trajectory modes were separated based on the time-resolved lMSD-analysis. The MSD curves were fitted using eq. 7 and 8, respectively. The resulting fitting parameters are summarized in Table S8.

**Figure S3:**
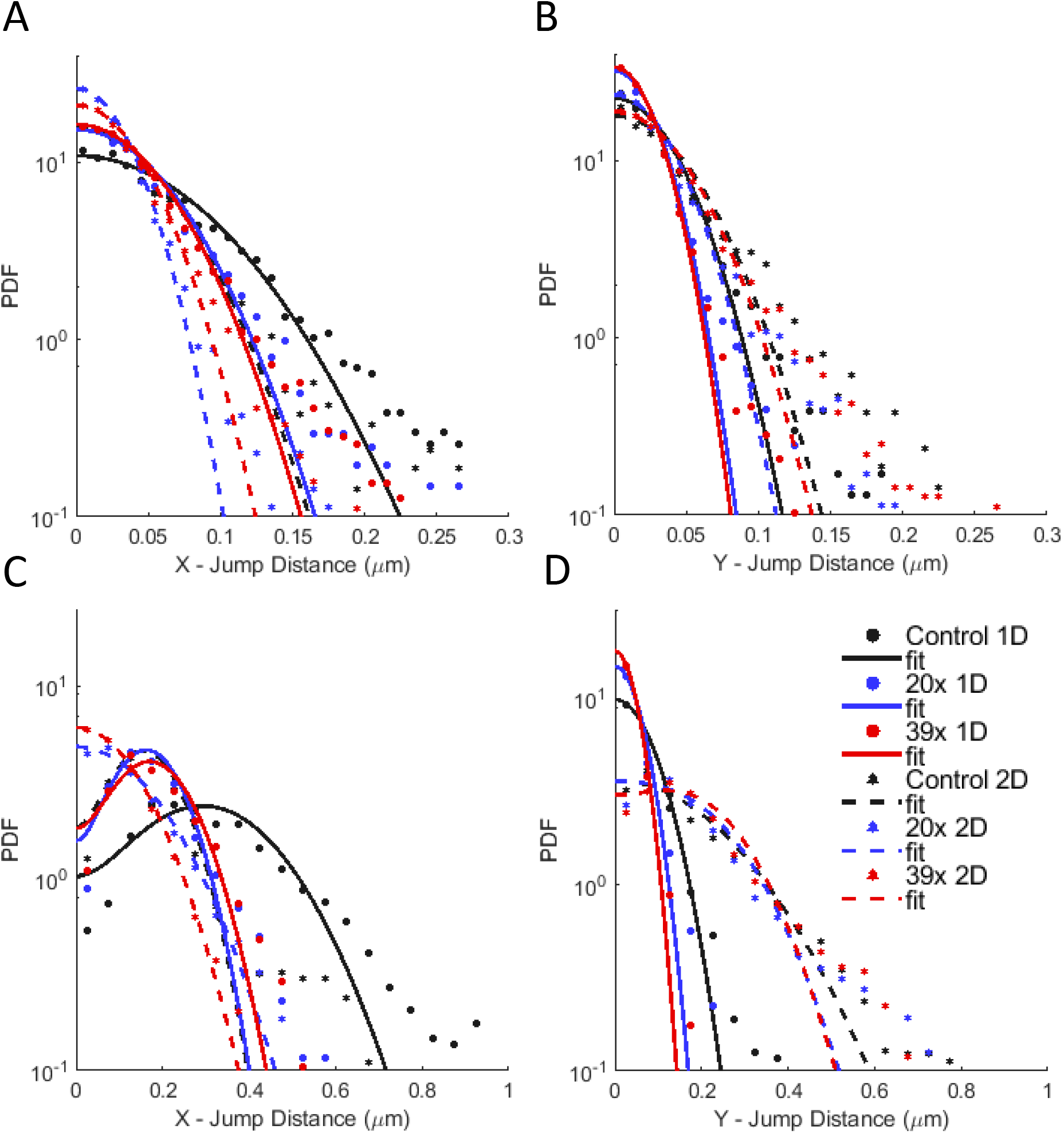
X- and Y- JDD PDFs of diffusive and super-diffusive trajectory parts. JDD PDFs calculated collectively for all diffusive trajectory parts along (A) the X- and (B) Y-axis. JDD PDFs calculated collectively for all super-diffusive trajectory parts along (C) the X- and (D) Y- axis. Color coding indicates data of lysosomes inside aligned (1D) and randomly oriented (2D) neurites of differentiated PC12 cells without (control) and with the G_4_C_2_ repeat expansion (20x and 39x repeats, respectively). Trajectory modes were separated based on the time-resolved lMSD-analysis. The JDD PDF curves were fitted using eq. 10 and 11, respectively. The resulting fitting parameters are summarized in Table S9.

